# *POL30* alleles in *Saccharomyces cerevisiae* reveal complexities of the cell cycle and ploidy on heterochromatin assembly

**DOI:** 10.1101/679829

**Authors:** Molly Brothers, Jasper Rine

## Abstract

In *Saccharomyces cerevisiae*, transcriptional silencing at *HML* and *HMR* maintains mating-type identity. The repressive chromatin structure at these loci is replicated every cell cycle and must be re-established quickly to prevent transcription of the genes at these loci. Mutations in a component of the replisome, the Proliferating Cell Nuclear Antigen (PCNA), encoded by *POL30*, cause a loss of transcriptional silencing at *HMR*. We used an assay that captures transient losses of silencing at *HML* and *HMR* to perform extended genetic analyses of the *pol30-6*, *pol30-8*, and *pol30-79* alleles. All three alleles destabilized silencing only transiently and only in cycling cells. Whereas *pol30-8* caused loss of silencing by disrupting the function of Chromatin Assembly Factor 1 (CAF-I), *pol30-6* and *pol30-79* acted through a separate genetic pathway but one still dependent on histone chaperones. Surprisingly, the silencing-loss phenotypes depended on ploidy but not on *POL30* dosage or mating-type identity. Separately from silencing loss, the *pol30-6* and *pol30-79* alleles also displayed high levels of mitotic recombination in diploids. These results established that histone trafficking involving PCNA at replication forks is crucial to the maintenance of chromatin state and genome stability during DNA replication. They also raised the possibility that increased ploidy may protect chromatin states when the replisome is perturbed.

## INTRODUCTION

Eukaryotic genomes include tightly packaged and transcriptionally repressed domains referred to as heterochromatin. The nucleosomes in heterochromatin are enriched for particular chromatin marks made by specialized chromatin-modifying enzymes. The marks left by these enzymes are recognized by other proteins that silence gene transcription. Although the exact histone modifications and heterochromatin proteins differ from organism to organism, there are unifying characteristics of heterochromatin including independence from underlying DNA sequence, replication late in S phase, and structural compaction.

To maintain the repression of genes within heterochromatin, histone modifications and chromatin-binding proteins must be faithfully replicated onto both daughter strands during DNA replication. The process that is required for inheritance of chromatin state through DNA replication is unclear but requires the interaction of chromatin regulators with various factors in the eukaryotic replisome (reviewed in Alabert and Groth 2017).

Proliferating Cell Nuclear Antigen (PCNA) is a DNA polymerase processivity clamp conserved from yeast to human (reviewed in Moldovan *et al.* 2017). PCNA is a homotrimer that assembles around individual DNA molecules, and, through protein-protein interactions, coordinates many activities at the DNA replication fork, including the processivity of DNA polymerase, Okazaki fragment processing, and chromatin assembly and remodeling. PCNA is also required for many different DNA repair pathways. Many chromatin modifiers and remodelers are recruited to replication forks through direct and indirect interactions with PCNA.

PCNA has a direct role in the stability of heterochromatin. In mice, heterochromatin Protein 1 (HP1) is recruited to replication forks through direct interaction with the histone chaperone complex Chromatin Assembly Factor 1 (CAF-I) (Murzina *et al.* 1999), which itself is recruited to replication forks through direct interaction with PCNA (Shibahara and Stillman 1999; Zhang *et al.* 2000; Ben-Shahar *et al.* 2009). Additionally, the maintenance of transcriptional silencing requires functional and stable DNA-bound PCNA in *Saccharomyces cerevisiae* (Zhang *et al.* 2000; Miller *et al.* 2008; Janke *et al.* 2018) These results suggest an important role for PCNA and CAF-I in the inheritance of chromatin states through DNA replication.

Circumstantial evidence for the importance of PCNA in the assembly of heterochromatin is also found in humans and *D. melanogaster.* In humans, Histone Deacetylase 1 (HDAC1), which is associated with transcriptional repression, interacts with PCNA *in vitro* and colocalizes with PCNA at replication forks *in vivo* (Milutinovic *et al.* 2002). In *D. melanogaster*, Polycomb Group (PcG) proteins, required for the establishment and maintenance of facultative heterochromatin, transiently associate with PCNA and CAF-I during DNA replication (Petruk *et al.* 2012).

*Saccharomyces cerevisiae* contains well-characterized heterochromatin domains that we used here to study the role of PCNA in epigenetic inheritance through DNA replication. Two of these loci, *HML* and *HMR*, share characteristics of heterochromatin in other organisms. Silencing of *HML* and *HMR* requires the activity of the SIR (<u>S</u>ilent Information <u>R</u>egulator) complex, composed of Sir2, Sir3, and Sir4. The Sir proteins are recruited first to the *E* and *I* silencers, nucleation sites flanking *HML* and *HMR*, and subsequently bind to nucleosomes that span the entire 3-4kb region between the silencers. Through the histone deacetylation activity of Sir2 and nucleosome-bridging ability of Sir3, the SIR complex creates a hypoacetylated, compact chromatin structure (reviewed in Gartenberg and Smith 2016).

In *S. cerevisiae*, alleles of PCNA, encoded by *POL30*, have been isolated that disrupt transcriptional silencing of reporter genes at telomeres and the silent mating-type locus, *HMR.* (Zhang *et al.* 2000). These alleles, *pol30-6*, *pol30-8*, and *pol30-79*, differ in phenotype and the degree of silencing loss they cause. Using the *ADE2* reporter at *HMR*, the *pol30-8* allele results in sectored colonies, suggesting the existence of two heritable states of gene expression: heritable silencing (*ADE2* expression off, resulting in red sectors), and heritable expression (*ADE2* expression on, resulting in white sectors). In contrast, colonies containing *pol30-6* or *pol30-79* are pink, suggesting a partial reduction of silencing in all cells (Zhang *et al.* 2000).

In combination with a deletion of *CAC1*, which encodes the large subunit of the histone chaperone CAF-I, the *pol30-6* and *pol30-79* alleles synergistically reduce silencing of *URA3* at telomere VII-L and of *ADE2* at *HMR*. However, the combination of *cac1*∆ and *pol30-8* result in similarly-sectored *ADE2* colonies as *pol30-8* alone and no further decrease in telomeric silencing than *pol30-8* alone. These two results suggest that PCNA may contribute to heritable silencing through at least two different mechanisms, one of which is through the histone-chaperone activity of CAF-I (Zhang *et al.* 2000).

Although reporter genes have a long history of successful use in genetic studies, the reliability of the *ADE2* and *URA3* reporters has been called into question, especially for situations involving DNA metabolism (Takahashi *et al.* 2011; Rossmann *et al.* 2011). Using a silencing-reporter assay that more sensitively captures loss-of-silencing events, better maintains the gene structure of *HML* and *HMR*, and is free of the complications of nucleotide metabolism, we have re-evaluated earlier claims about the silencing phenotypes of *pol30-6*, *pol30-8*, and *pol30-79*, extended the analyses substantially, and have provided new interpretations of published observations.

## MATERIALS AND METHODS

### Yeast Strains

All strains in this study were derived from W303 and are listed in Table S1. Plasmids used in this study are listed in Table S2. Gene deletions (except for *bar1*∆) were created by one-step integration of PCR-amplified disruption cassettes (Goldstein *et al* 1999; Güldener *et al.* 2002), using primers listed in Table S3. The *pol30-8* (R61A, D63A) allele, *mcm2-3A* (Y79A, Y82A, Y91A) allele, and *bar1*∆ were introduced using Cas9 technology. Guide RNAs targeting *POL30*, *MCM2*, and *BAR1* are listed in Table S3. The sgRNA dropout-Cas9 expression plasmid (pJR3428) was assembled using a toolkit from Lee *et al.* 2015. The gRNA target and non-target strands were integrated into pJR3428 by Golden Gate cloning using the restriction enzyme *Bsm*BI as described in Lee *et al.* 2015. The repair templates were made by annealing oligos in Table S3 and extending the 3’ ends using Phusion Polymerase (New England Biolabs). The *pol30-6* (D41A, D42A) and *pol30-79* (L126A, I128A) alleles were created by integrating gene blocks containing each allele along with the selectable marker *URA3*, found in Table S3. A detailed description of this method is in Supplemental Information File S1. Double mutants were created by genetic crosses and were confirmed by tetrad analysis, using selectable markers for gene disruptions and mutant-specific PCR and/or sequencing for *pol30-6*, *pol30-8*, *pol30-79*, and *mcm2-3A*. Creation of *POL30* hemizygotes and the tetraploid strain (JRY12026) used plasmid shuffles with pBL230-0 [*CEN/ARS TRP1 POL30*] (Ayyagari *et al.* 1995; Zhang *et al.* 2000), described in detail in Supplementary Information File S1.

### Colony Growth and Imaging

Strains were first patched onto selective medium plates to select for cells expressing hphMX, and thus had not lost silencing and excised the RFP-hphMX cassette (Figure 1A): YPD containing 200μg/mL G418 (Geneticin; Life Technologies) for strains carrying the kanMX cassette or YPD containing 300μg/mL Hygromycin B (MilliporeSigma) for strains carrying the hphMX cassette. Cells were then resuspended in water and plated onto Complete Supplement Mixture (CSM) or CSM – Trp (Sunrise Science Products), 1.5% agar, at a density of ~30 cells per plate. Colonies were imaged after 5-6 days of growth at 30°C. At least 10 colonies per genotype were imaged using a Leica M205FA fluorescence stereomicroscope, a Leica DFC3000G CCD camera, and a PlanApo 0.63X Objective. Image analysis and assembly was performed using ImageJ software (Fiji, Schindelin *et al.* 2012).

**Figure 1.**
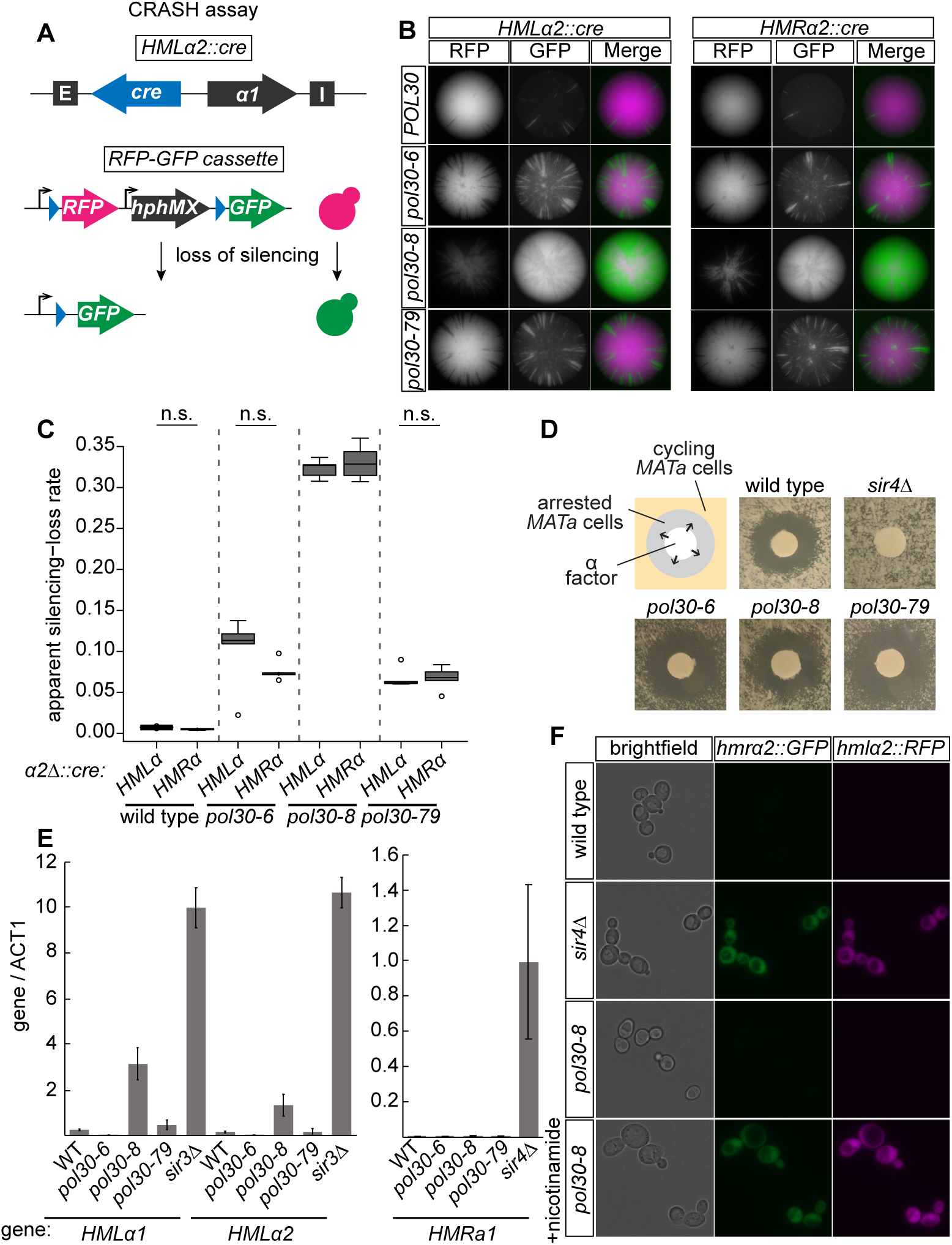
Alleles of *POL30* caused transient loss of silencing. **(A)** Schematic of the CRASH (Cre-Reported Altered States of Heterochromatin) loss-of-silenc­ ing assay. Expression of *ere* from *HMLa2::cre* occurs when transcriptional silencing is disturbed. In cells that lose silencing even transiently, Cre causes a permanent switch from expressing RFP to expressing GFP. In a similar strain, ere is expressed from *HMRa2* to detect loss-of-silencing events at HMR. **(B)** Colonies of *HMLa2::cre* (left panel) and *HMRa2::cre* (right panel) strains for each *POL30* allele. Each green sector represents a loss-of-silencing event. Wild-type strains (JRY10790, left and JRY10710, right) had few sectors. Strains containing *po/30-6* (JRY11137, *po/30-6po/30-8po/30-79* left and JRY11186, right}, *po/30-8 (JRY11188, left and JRY11187, right}*, or *po/30 -7 9* (JRY11141, left and JRY11608, right) had elevated sectoring compared to wild type. **(C)** The apparent silencing-loss rates for each of the strains in (B) were quantified by flow cytometry as described in Materials & Methods and in Janke *et al.* 2018. Significant differences (*) or not-significant differences (n.s.) were determined by one-way ANOVA and Tukey’s Honestly Signficant Difference (HSD) post-hoc test. The center line of each box plot represents the median of at least 5 biological replicates. The boxes represent the 25th and 75th percentiles. Whiskers represent the range of values within 1.5X the interquar­ tile range. Values extending past 1.5X the interquartile range are marked as outliers (circles). **(D)** a-factor halo assay. Filter papers soaked in the mating pheromone a-factor (200µM in 100mM sodium acetate) were placed onto a freshly-spread lawn of *MATa* cells of each indicated genotype. *MATa* cells that maintain silencing at *HMLa* will arrest in G1 phase around the filter paper, creating a ’halo’. Cells that heritably lose silencing at *HMLa* do not arrest in response to a-factor. Representative images of wild type (JRY4012), *sir4t,.* (J RY4577}, *po/30-6* (JRY11645 }, *po/30-8* (JRY11647}, and *po/30-79* (JRY11649) are shown. **(E)** RT-qPCR of *a 1* and *a2* transcripts from *HMLa* and *a1* from *HMRa.* Quantification was performed using a standard curve for each set of primers and normalized to *ACT1* transcript levels. Error bars represent standard deviation. Bars represent the normalized average of three technical replicates of each indicated strain: WT (JRY11699 *matt,.), sir3t,.* (J RY9624, *matM mrt,.) sir4t,.* (JRY12174 *MATa), po/30-6* (JRY11700 *mat* t,.), *po/30-8* (JRY11701 maM), *po/30-79* (JRY11702 *matt,.).* **(F)** Genes encoding fluorescent reporters were placed at *HMLa2* (RFP) and *HMRa2* (GFP) reported on transcription from the two loci at the same time with single-cell resolution. Representative images of each strain are shown below the schematic: WT (wild type, JRY11129), *sir4t,.* (JRY11131), *po/30-8* (JRY11132). The last row is the *po/30-8* strain (JRY11132) treated with 5mM nicotinamide, a SIR2 inhibitor, prior to imaging.

### Quantification of Silencing Loss by Flow Cytometry

For each strain, 3-5 single colonies were inoculated in 1mL in selective media to select for cells that had not lost silencing. These 1mL cultures were grown in deep 96-well plates (VWR) at 30°C overnight to saturation. Overnight cultures were diluted into 1mL of fresh, non-selective YPD to ~10^5^ cells/mL in a deep 96-well plate and grown for 5-6 hours before flow cytometry. For each culture, a minimum of 15,000 events were collected using a BD High-Throughput Sampler on a BD LSR Fortessa X20 Cell Analyzer. Gating and quantification were performed as previously described (Janke *et al.* 2018).

### α-factor Halo Assay

*MATa* strains were scraped from a YPD plate with a toothpick, resuspended in water, and ~150,000 cells were freshly spread onto Complete Supplement Mixture plates (CSM, Sunrise Science Products). Hole-punched Whatman filter papers were soaked for ~5 sec in 200μg/uL α-factor in 100mM sodium acetate and then placed onto the plates. Three soaked filter-paper circles were placed on each plate. Plates were incubated at 30°C for 36-48 hours before imaging.

### RNA preparation for quantitative qPCR

Cells were grown to mid-log phase in YPD and RNA was extracted using hot acidic phenol and chloroform (Collart and Oliviero 2001). Samples were treated with DNase I (New England Biolabs) and subsequently purified using the Qiagen RNeasy Mini Kit. Complementary DNA (cDNA) was synthesized using the SuperScript III First-Strand Synthesis System (Invitrogen) and oligo(dT) primers. Quantitative PCR of cDNA was performed using the DyNAmo HS SYBR Green kit (Thermo Fisher Scientific) on an Mx3000P machine (Stratagene) using the primers listed in Table S3. Standard curves were generated using the *sir3*∆ *mat*∆ *hmr*∆ (JRY9624) or *sir4*∆ strain (JRY12714).

### Live-cell Imaging

All single-cell microscopy images were collected on a Zeiss Axio Observer Z1 inverted microscope equipped with a Plan-Apochromat 63x oil-immersion objective (Zeiss) and the Definite Focus System for maintenance of focus over time. yEGFP was excited with the 420-500nm spectrum range at 20% intensity, and yEmRFP was excited with the 500-755nm spectrum range at 20% intensity from a CoolLED pE-300 ultra and collected with the Multiband Semrock Filter (LF405/488/594-A-ZHE).

Images were acquired with a Teledyne Photometrics Prime 95B sCMOS camera. For time-lapse experiments, images were collected every 5 or 7 minutes, using an exposure time of 20msec for brightfield, 50msec for yEGFP, and 200msec for yEmRFP. At each time point, multiple stage positions were collected using an ASI MS-2000 XYZ piezo stage. The microscope, camera, and stage were controlled with the Micro-manager software (Edelstein *et al.* 2014). Image analysis and assembly was performed using Fiji software (Schindelin *et al.* 2012).

For the CRASH time course setup, each strain was inoculated in 5mL of selective media to select for cells that had not lost silencing. Cultures were grown to saturation overnight at 30°C. Overnight cultures were diluted back to 10^6^ cells/mL in 10mL of YPD containing G418 or Hygromycin B and grown to early-log phase (~ 4 × 10^6^ cells/mL). The culture was then split into 5mL of YPD containing G418 or Hygromycin B and 40nM α-factor in 100mM sodium acetate, and 5mL of YPD containing G418 or Hygromycin B. The cultures were grown for another 90 minutes (~1 doubling). The cells were then harvested and resuspended in water. The resuspended cells were diluted to ~2 × 10^7^ cells/mL, and 5uL were pipetted onto a 1cm x 1cm square of CSM – Trp + 2% agar with or without 40nM α-factor in 100mM sodium acetate. Both agar pads were placed into a 27mm glass dish (Thermo Scientific) and mounted in a Pecon Incubator XL with Heating Unit XL S (Zeiss) controlled by TempModule S (Zeiss) and kept at 30°C for the duration of the experiment. To calculate the number of switches per 10,000 cells in the arrested condition, the number of cells that were RFP-expressing at time zero were counted. The total number of switches (RFP-to-GFP) were counted for all of those cells and divided by the total number of RFP-expressing cells at time zero. To calculate the number of switches per 10,000 divisions in the cycling condition, the number of switches over the entire time course was divided by the calculated total number of divisions. The number of divisions was calculated using the following formula:

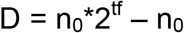

value t: total number of minutes in the time course (480 minutes for an 8hr time course) value f: division rate (per minute). To determine this value, the time from small bud to the next small bud for five cells at the beginning of the time course, five cells at the end of the time course in the center of a micro-colony, and five cells at the end of the time course at the edge of a micro-colony was averaged to get the time for one division. value n_0_: number of RFP-expressing cells at time 0

### Protein Isolation and Immunoblotting

Each strain was inoculated in 5mL of YPD and grown overnight to saturation. Overnight cultures were diluted to ~2 × 10^5^ cells/mL in fresh YPD and grown to mid-log phase, and ~10^8^ cells were harvested and pelleted. Pellets were resuspended in 1mL of 5% trichloroacetic acid and incubated at 4°C for 90 minutes. The precipitates were pelleted, washed twice with 1mL of 100% acetone, and air-dried. Dried pellets were resuspended in 100uL of protein breakage buffer (50mM Tris-HCl pH 7.5, 1mM EDTA, 3mM DTT) and an equal volume of 0.5mm zirconium ceramic beads (BioSpec Products) followed by five cycles of vortexing, 1min bursts with 1min of incubation on ice between each cycle. 50μL of 3X SDS sample buffer (188mM Tris-HCl pH 6.8, 30% glycerol, 150mM DTT, 6% SDS, 0.03% bromophenol blue, 2% BME) was added to each sample and incubated at 95°C for 5min. Insoluble material was pelleted by centrifugation and an equal volume of the soluble fraction from each sample was run on an SDS-Polyacrylamide gel (Mini-PROTEAN TGX Any kD pre-cast gel, BioRad) and transferred to a nitrocellulose membrane using a TransBlot Turbo Transfer Pack (BioRad) on the Mixed MW setting of a TransBlot Turbo machine (BioRad). The membrane was cut horizontally in half between the 50kD and 37kD markers and separated to blot for Hxk2 on the top half and PCNA on the bottom half. The membranes were blocked in Odyssey Blocking Buffer (LI-COR Biosciences), and the following primary antibodies and dilutions were used for detection: PCNA (Abcam ab221196, 1:1000), Hxk2 (Rockland #100-4159, 1:10,000). The secondary antibody used was IRDyeCW800 goat anti-rabbit (LI-COR Biosciences, 1:20,000), and the membrane was imaged on a Li-Cor Odyssey Imager. All washing steps were performed with Phosphate Buffered Saline + 0.1% Tween-20. Quantitative analysis was performed using Fiji software (Schindelin *et al.* 2012): The area under the intensity peak above background for each band was used for normalization to Hxk2 followed by comparison between lanes.

### Tetrad Analysis

Diploid cells were sporulated on 1% potassium acetate, 2% agar, 0.25X CSM plates for 2-3 days at room temperature. Tetrads were dissected onto YPD plates using a micromanipulator and grown for 2 days before replica plating and scoring.

### Patch Mating Assay

All strains were patched on YPD and grown at 30°C for 3 days. A small sample of yeast was scraped from the center of each patch and patched onto a fresh YPD plate and grown overnight at 30°C. The following day, the YPD plate was replica plated onto a YPD plate with a fresh lawn of *MATa* haploid testers and a YPD plate with a fresh lawn of *MAT*α haploid testers. The replica plates were grown overnight and then replica plated onto minimal media plates the next day. Minimal media plates were grown for 2 days at 30°C and then imaged.

### Data Availability

All data necessary for confirming the conclusions presented in the article are represented fully within the article. Supplemental material has been uploaded to the GSA figshare public repository. File S1 contains supplementary materials and methods. Table S1 contains details about the yeast strains used in this study. Table S2 contains the plasmids used in this study. Table S3 contains oligonucleotides used in this study.

## RESULTS

### Alleles of *POL30* caused transient loss of silencing

We introduced alleles of *POL30* implicated in heterochromatic silencing, *pol30-6, pol30-8*, and *pol30-79* (Zhang *et al. 2000*) into a strain we previously constructed that allows sensitive detection of losses of heterochromatin silencing (Figure 1A, Dodson and Rine 2015). In this strain, the α*2* coding sequence at *HML*α or *HMR*α is replaced with the coding sequence of Cre recombinase. The *URA3* locus on chromosome V is replaced by *loxP* sites flanking the *RFP* gene and the selectable marker *hphMX* downstream of the strong *TDH3* promoter. Downstream of *loxP-RFP-hphMX-loxP* is a promoter-less *GFP* gene. Upon loss of silencing at *HML*α or *HMR*α, Cre recombinase is expressed and excises the *RFP* and *hphMX* sequences, resulting in a permanent switch from expressing RFP to expressing GFP (Figure 1A). Within a colony, a sector of green cells represents a loss-of-silencing event in a cell born at the vertex with the sector representing growth of the descendants following the loss event. This assay is referred to as the CRASH assay (<u>C</u>re-<u>R</u>eported <u>A</u>ltered <u>S</u>tates of <u>H</u>eterochromatin, Dodson and Rine 2015).

Each of the *pol30* mutants resulted in increased sectoring compared to wild-type *POL30* at both *HML*α and *HMR*α (Figure 1B). We also quantified loss-of-silencing events in these strains using flow cytometry. The apparent silencing-loss rate was calculated as the number of yellow cells (cells that had recently excised the *RFP* gene) divided by the sum of all yellow cells *and* red cells. The loss rates from flow cytometry experiments mirrored qualitative assessments of loss rates from colony sectoring (Figure 1C). *pol30-8* cells had the most unstable silencing followed by *pol30-6* and then *pol30-79* (Figure 1B, 1C).

The CRASH assay reveals how unstable transcriptional silencing is in a given strain, but because it is a permanent switch, the assay is unable to capture the heritability of the de-repressed state at *HML* and *HMR*. To determine how heritable the loss-of-silencing events were in strains with the *pol30-6*, *pol30-8*, and *pol30-79* alleles, we first performed an α-factor halo assay (Figure 1D). When *MATa* cells are exposed to the mating pheromone α-factor, they arrest in G1. On a lawn of *MATa* cells, this results in a halo of arrested cells surrounding the source of α-factor (Figure 1D, wild type) However, if *MATa* cells lose silencing at *HML*α, they no longer arrest in response to α-factor (Figure 1D, *sir4*∆). If the loss-of-silencing events created by the *pol30* mutants were heritable, colonies would grow within the halo, as noted for the *sir4*∆ strain. However, for all three alleles, we observed no cell growth within the halos (Figure 1D).

In agreement with the α-factor halo results, strains containing *pol30-6, pol30-8*, or *pol30-79* also showed only low levels of *HML*α*1*, *HML*α*2*, and *HMRa1* transcripts by RT-qPCR (Figure 1E). Analysis of *cre* transcripts from CRASH strains with the *POL30* alleles also revealed only low levels of transcription (data not shown).

The absence of a notable increase in transcripts from *HML* and *HMR* was particularly surprising for *pol30-8*, which had an extremely high sectoring rate (Figure 1B, 1C) and was previously suggested to have bi-stable epigenetic states based upon the *HMR::ADE2* reporter (Zhang *et al.* 2000; Miller *et al.* 2010). Therefore, we also placed this allele in another reporter strain that encodes *GFP* at *HMR*α*2* and *RFP* at *HML*α*2*. If the *pol30-8* allele resulted in a population of cells with stable expression from *HML* or *HMR*, we would expect some cells to be fluorescent. Instead, *pol30-8* cells with these reporter loci expressed no detectable fluorescence (Figure 1F). Importantly, *GFP* and *RFP* were expressed in this strain when silencing was lost following treatment with a *SIR2* inhibitor, nicotinamide (Figure 1F).

### Loss-of-silencing events in strains with defective *POL30* alleles occurred predominantly in cycling cells

Given the major role of PCNA in DNA replication, we considered that the loss-of-silencing events may occur only during S-phase. Alternatively, because PCNA is involved in replication-independent roles such as DNA repair, it was possible that heterochromatin assembled in the *pol30* mutants might be unstable at any point in the cell cycle. We performed time-lapse microscopy using CRASH strains to compare the rate that silencing is lost in G1-arrested cells to the rate in cycling cells. As an example, in cycling *pol30-8* cells, switches were readily visible over the time-course of 8 hours (Figure 2A). In wild-type cells, the rate of switching was about the same for arrested versus cycling cells. However, in cells containing each of the *pol30* mutants, losses of silencing predominantly occurred in cycling cells. Arrested *pol30* mutants exhibited a low frequency of silencing loss comparable to that seen in wild type (Table 1). These results suggested that the *pol30-6*, *pol30-8*, and *pol30-79* alleles caused only transient losses of silencing in actively cycling cells, with quick re-establishment of the silent state.

**Table 1.**
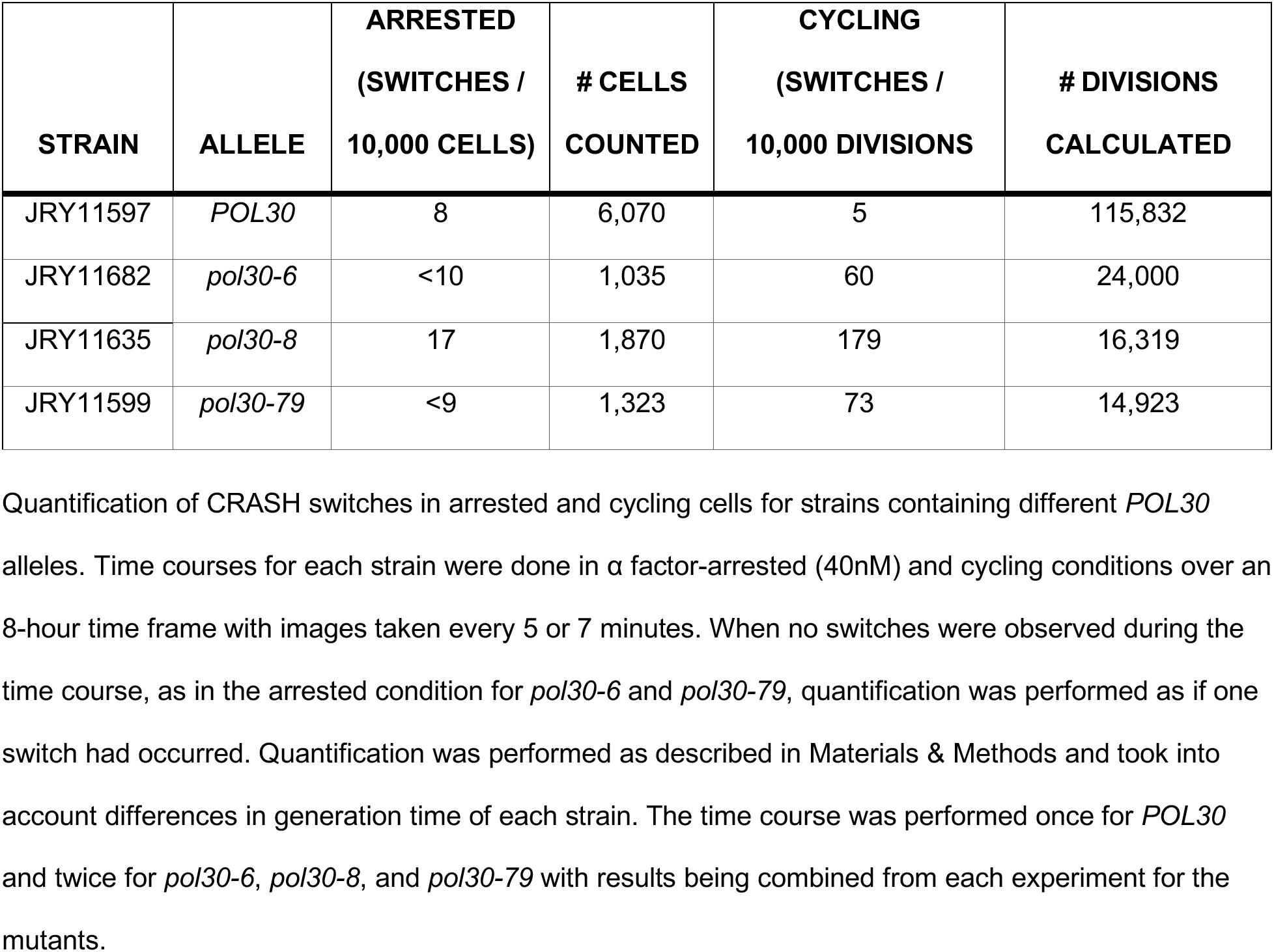
Loss-of-silencing events in strains with defective *POL30* alleles occurred predominantly in cycling cells

**Figure 2.**
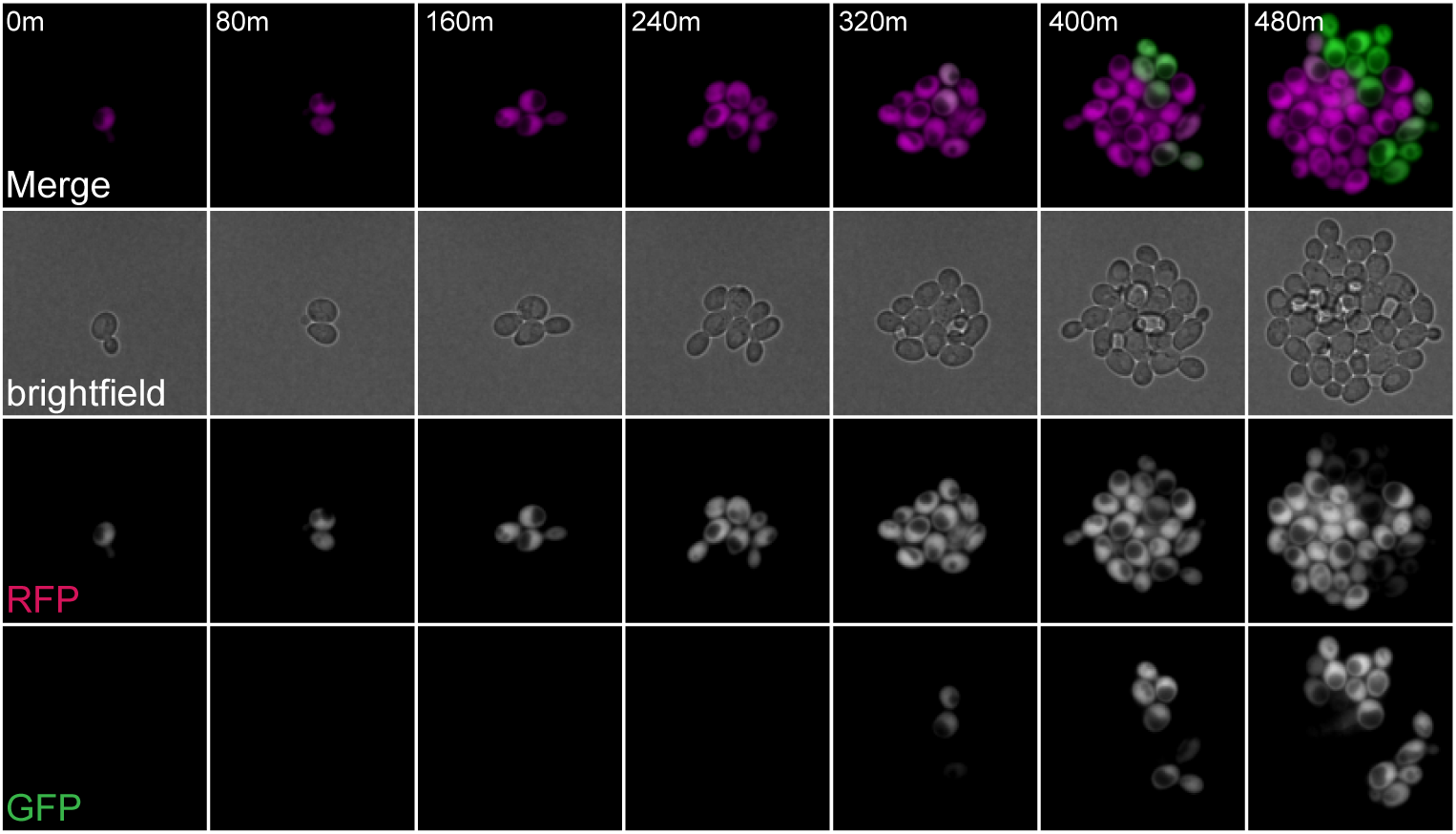
Loss-of-silencing events in strains with defective *POL30* alleles occurred predominantly in cycling cells. Representative images of 2 loss-of-silencing events in one micro-clone of cycling cells in the *HMLa2::cre* CRASH assay containing the *po/30-8* allele (JRY11635, *bar1Δ*). See Table 1 for calculation of switching rates for all *POL3O* alleles.

### Silencing loss caused by *POL30* alleles was dependent on ploidy

To determine whether each of the *pol30* mutants disrupted silencing in the CRASH assay through the same mechanism, we performed pairwise complementation testing among the three alleles. As a necessary prerequisite, we tested each allele in a diploid in combination with wild-type *POL30. pol30-6, pol30-8*, and *pol30-79* were all recessive to *POL30* by this assay (Figure 3A).

**Figure 3.**
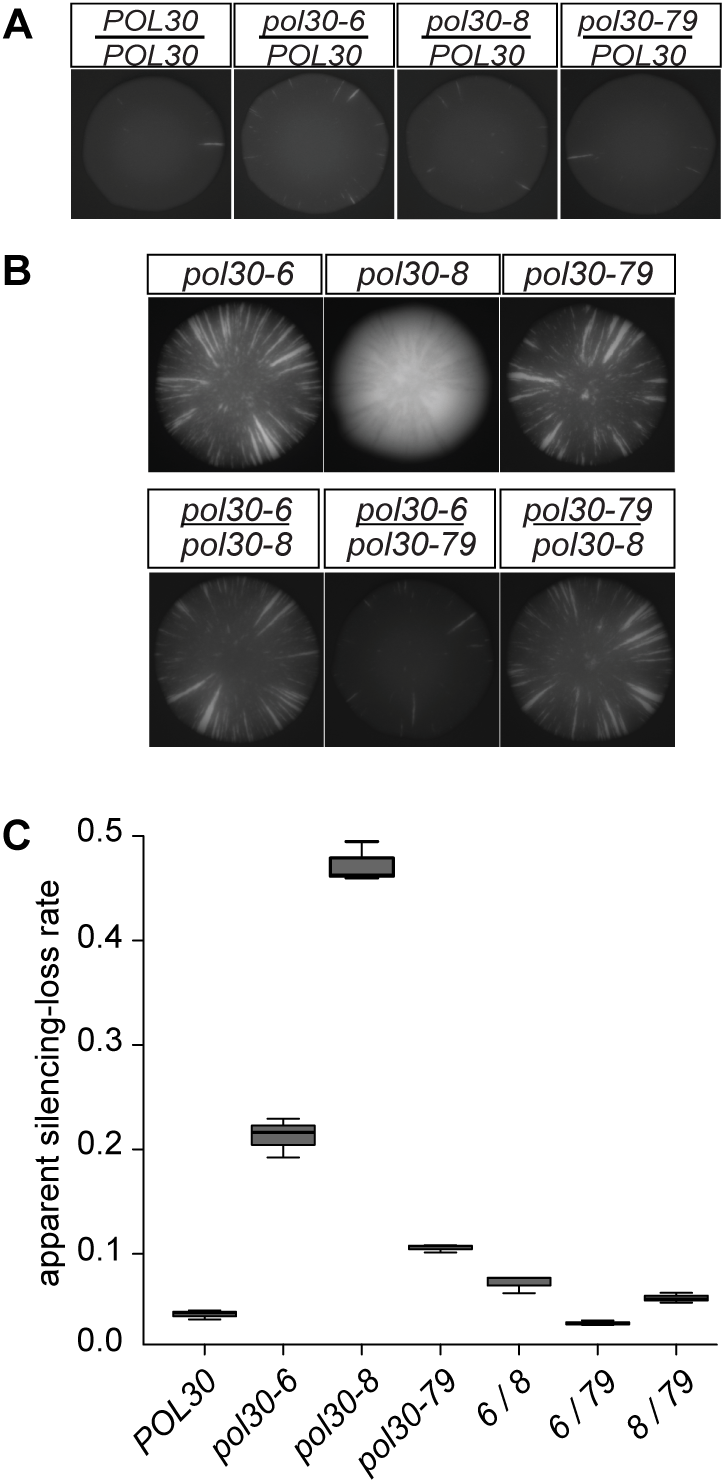
*POL30* alleles complemented in diploids but not in haploids. **(A)** Each *po/30* allele was recessive to wild-type *POL30* in the CRASH assay with ere at *HMLa2: POL30/POL30* (JRY11159), *po/30-6/POL30* (JRY11160), *po/30-8/POL30* (JRY11169), *po/30-79/POL30* (JRY11161). Only the GFP channel is shown. These diploid strains contained only one *HMLa2::cre* and one *RFP-hphMX-GFP* cassette. **(B)** Complementation of *po/3 0-6, po/30-8*, and *po/30-79* in the CRASH assay with ere at *HMLa2*. Only the GFP channel is shown. The top row shows representative haploid colonies containing the indicated allele: *po/30-6* (JRY11137), *po/30-8* (JRY11188), *po/30-79* (JRY11141). The bottom row shows representative diploid colonies containing a combination of the indicated alleles: *po/30-6/po/30-8* (JRY11656), *po/30-6/po/30-79* (JRY11657), *po/30-Blpo/30-79* (JRY11658). Diploid strains contained only one *HMLa 2::cre* and one *RFP-hphMX-GFP* cassette. **(C)** The apparent silencing-loss rates for each of the strains in (B) and *POL30* (JRY10790) were quantified by flow cytometry as described in Figure 1C.

If two recessive *pol30* mutants disrupt heterochromatin through different mechanisms, then the combination of those two alleles in the diploid should complement, decreasing the frequency of RFP-to-GFP switches compared to each allele alone. All three combinations of *pol30* mutants in heteroallelic diploids decreased sectoring relative to haploids with each allele individually, most dramatically evident in the *pol30-6 / pol30-8* diploid (Figure 3B, 3C). Because the sectoring phenotype of *pol30-79* was weak on its own, its effect in combination with the other alleles was not as striking but still noticeably in combination with both *pol30-6* and *pol30-8* (Figure 3B, 3C).

In the simplest manifestation of the complementation test in yeast genetics, the phenotype of haploids containing each mutant of interest is compared to the phenotype of diploids containing both mutations, often ignoring potential complications of ploidy in assessing whether the mutations complement. Surprisingly, homozygosity of each allele at least partially suppressed the loss-of-silencing phenotype as measured by the CRASH assay (Figure 4A, 4B, 4C).

**Figure 4:**
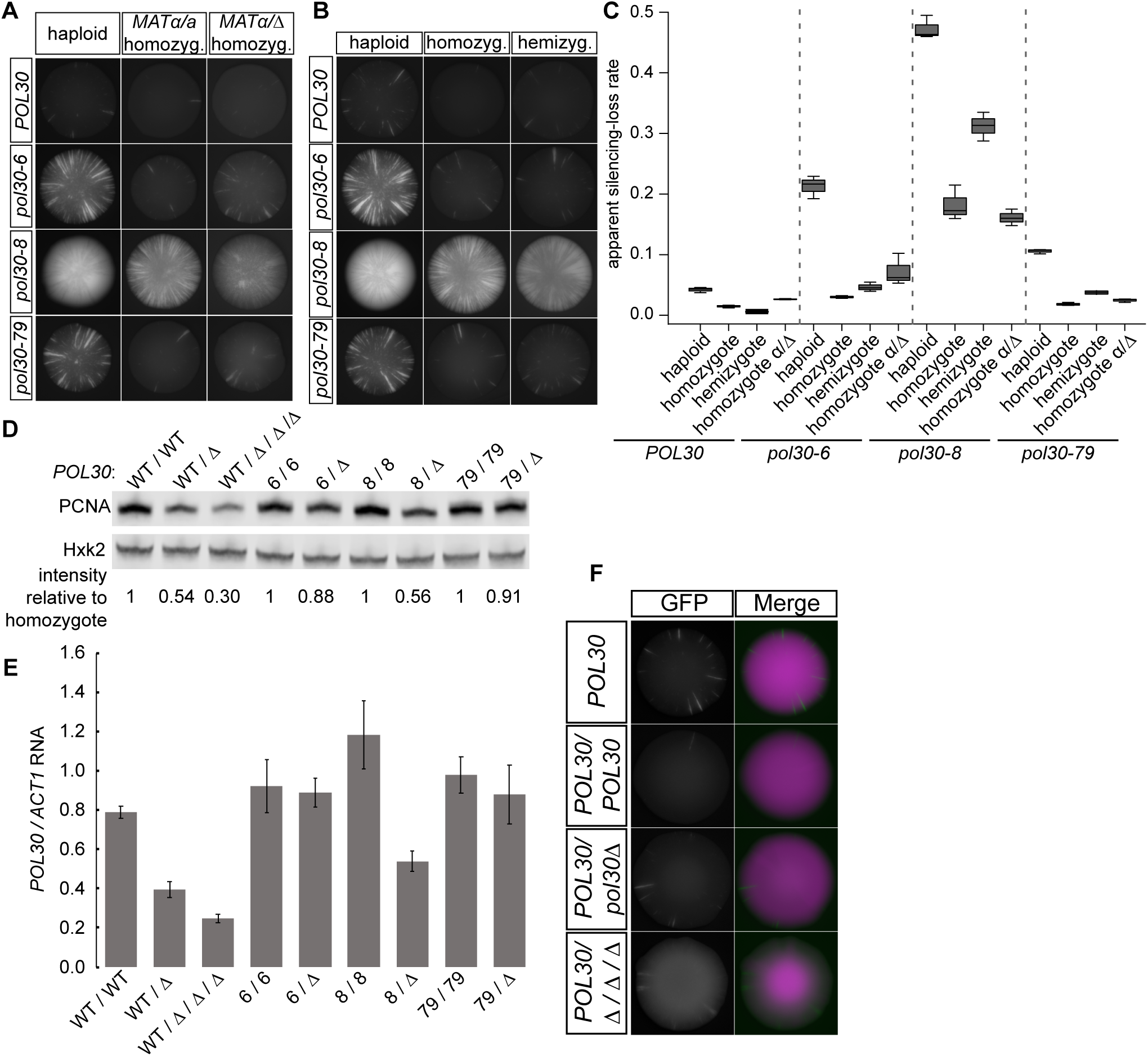
The effect of *POL30* alleles on silencing was dependent on ploidy. **(A)** Representative images of haploids, *MATa/MATa* homozy­ gotes, and *MATalmatt:.* homozy­ gotes for each indicated *POL30* allele in the CRASH assay with ere at *HMLa2.* Homozygous diploid strains contained two copies of each indicated allele. Only the GFP channel is shown. Diploid strains contained only one *HMLa 2::cre* and one *RFP-hphMX-GFP* cassette. *POL30* row: JRY10790, JRY11159, JRY11718. *po /30 -6* row: JRY11137, JRY11686, JRY11719. *po/30-8* row: JRY11188, JRY11687, JRY11744. *po/30-79* row: JRY11141, JRY11688, JRY11720 **(B)** Representative images of haploids, homozygotes, and hemizygotes for each indicated *POL30* allele in the CRASH assay with ere at *HMLa2.* Homozygotes are diploid strains containing two copies of each indicated allele. Hemizygotes are diploid strains containing one copy of the indicated allele over a deletion of *POL30 (po/30 !::.).* Diploid strains contained only one *HMLa2::cre* and one *RFP-hphMX-GFP* cassette. *POL30* row: JRY10790, JRY11159, JRY11745. *po/30-6* row: JRY11137, JRY11686, JRY11822. *po/30-8* row: JRY11188, JRY11687, JRY11749. *po/30-79* row: JRY11141, JRY11688, JRY11823. **(C)** The apparent silencing-loss rates for each of the strains in (A) and (B) were quantified by flow cytometry as described in Figure 1C. **(D)** lmmunoblot analysis of PCNA protein levels in homozygotes and hemizygotes of each allele (same strains as Figure 4B) as well as a tetraploid containing just one copy of wild-type *POL30 (WTI MM t:.*, J RY12026). The tetraploid contains two copies of *HMLa2::cre* and the *RFP-hphMX-GFP* cassette. Hxk2 levels served as a loading control. *POL30* allele nomenclature was abbreviated. Each PCNA band intensity was normalized to Hxk2 intensity. After normalization to Hxk2, the relative intensity of each lane to its corresponding *POL30, po/30 −6, po /30 -8*, or *po/30-79* homozygote was calculated and displayed. **(E)** RT-qPCR analysis of *POL30* RNA levels in homozygotes and hemizygotes of each allele and a tetraploid with one copy of wild-type *POL30* (same strains as Figure 4B and 4D). Error bars represent standard deviation. Quantification was performed using a standard curve for each set of primers and normalized to *ACT1* transcript levels. Error bars represent standard deviation. Bars represent the normalized average of three technical replicates of each indicated strain. **(F)** Representative images of a wild-type haploid *(POL30* JRY10790), homozygote *(POL30/POL30* JRY11159), hemizygote *(POL30/t:.* JR Y11745), and tetraploid with one copy of POL30 *(POL30/MMt:.* JRY12026). The haploids and diploids contained only one *HMLa2::cre* and one *RFP-hphMX-GFP* cassette. The tetraploid contained two copies of *HMLa2::cre* and the *RFP-hphMX-GFP* cassette. The increased background in the GFP channel of the tetraploid was due to loop-out of one *RFP-hphMX* cassette, leaving just one *RFP-hphMX-GFP* cassette able to switch.

To test whether the phenotypic suppression of *pol30* mutations reflected mating-type differences between haploids and diploids, we created *MAT*α */ mat*∆ diploids homozygous for each *POL30* allele. These cells, though diploid, express only α-specific genes and therefore behave as *MAT*α haploids. If mating-type were the cause of sectoring suppression in the *pol30-6, pol30-8, and pol30-79* diploids, then *MAT*α */ mat*∆ diploids would be expected to increase the sectoring rate back to the same level as the haploids. For all three alleles, changing the mating type had little or no effect on the reduced sectoring phenotype of diploids (Figure 4A, 4C). Therefore, mating type was not responsible for the difference between haploid and diploid *pol30* mutants.

Alternatively, the reduced sectoring in diploids could reflect a difference in gene dosage of *POL30* between haploids and diploids. We therefore created hemizygotes for each allele in which diploids contained only one copy of the allele instead of two. Although hemizygosity did not increase the sectoring rate to the same level as the haploid, the *pol30-8* hemizygote had a statistically significant increase in the loss of silencing rate compared to the homozygote (Figure 4C). In contrast to *pol30-8*, the *pol30-6* and *pol30-79* hemizygotes had only minor, statistically-insignificant increases in sectoring (Figure 4B, 4C). Immunoblot of Pol30 protein levels and qRT-PCR of *POL30* RNA levels in the various mutants revealed that *pol30-6* and *pol30-79* expression was comparable in hemizygotes and homozygotes, whereas wild-type *POL30* and *pol30-8* expression decreased by half at both the protein and RNA level (Figure 4D, 4E).

The effect of *pol30* mutants on heterochromatin stability was different in haploids and diploids, and this difference was independent of mating type and largely independent of gene dosage. Moreover, silencing in tetraploid cells with just one copy of *POL30* was as stable in haploids and homozygote or hemizygote diploids (Figure 4F), even with a quarter the expression of *POL30* relative to the amount of chromatin the cell (Figure 4D, 4E).

### *pol30-6* and *pol30-79* caused high rates of mitotic recombination and gene conversion in diploids

In the course of characterizing the *POL30* mutants, we found that *pol30-6* and *pol30-79* homozygous diploids had high rates of mitotic recombination/gene conversion in diploids that was dependent on mating type but not on *POL30* gene dosage.

Completely green CRASH colonies of *pol30-6* and *pol30-79* homozygotes revealed mitotic recombination of the *loxP-GFP* cassette through the existence of sectors that were twice as bright and sectors that were non-fluorescent suggesting duplication and loss of the *GFP* cassette, respectively (Figure 5A). *pol30-8* homozygotes did not display mitotic recombination in completely green colonies (data not shown). In *mat*∆ */ MAT*α diploids homozygous for *pol30-6* and *pol30-79*, this phenotype was suppressed (Figure 5A). Hemizygosity for *pol30-6* or *pol30-79* did not suppress the high levels of *loxP-GFP* mitotic recombination seen in homozygous diploids (data not shown).

**Figure 5:**
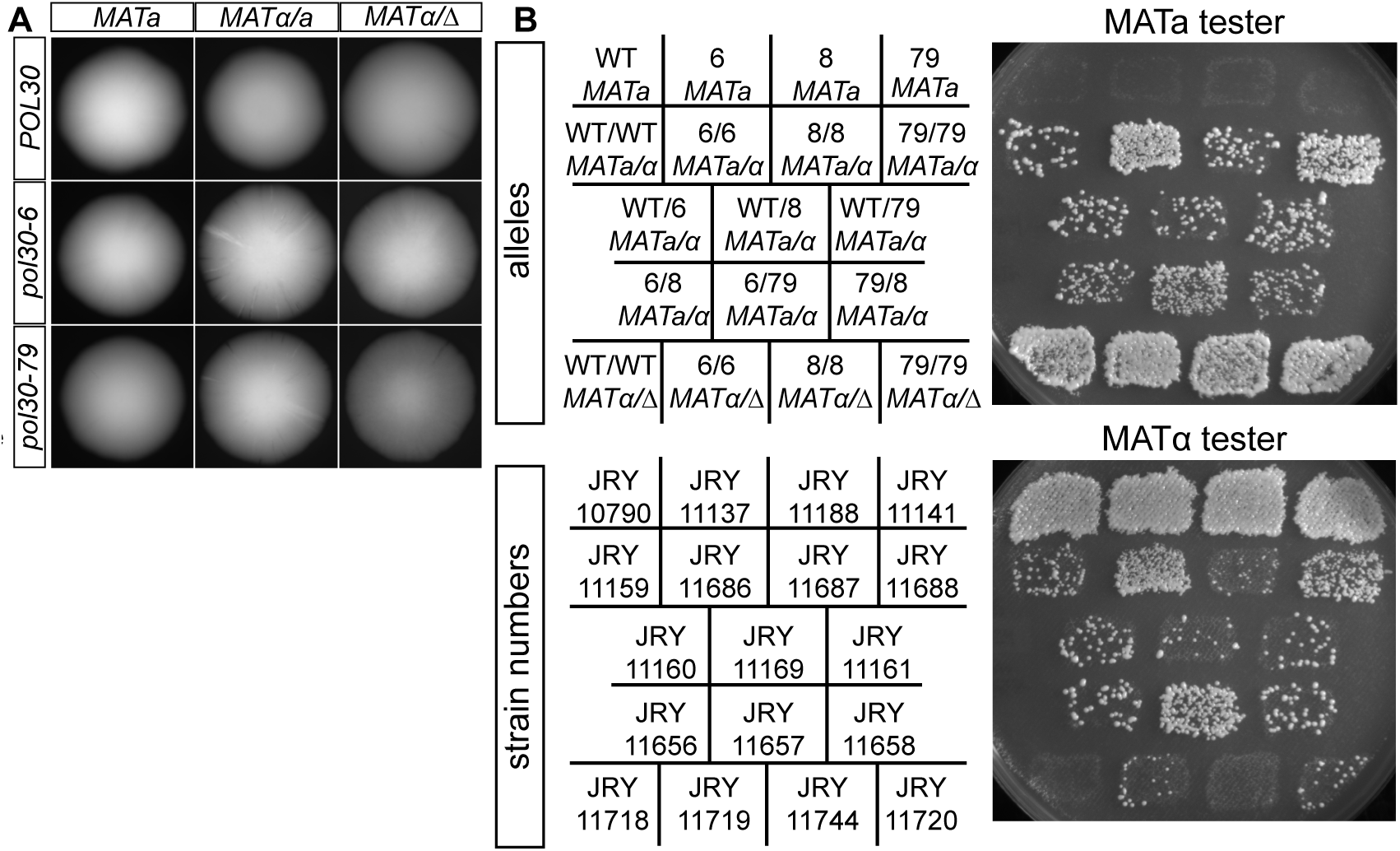
*po/30-6* and *po/30-79* caused high rates of mitotic recombination and gene conversion in diploids. Representative colonies of each indicated genotype that were completely green from the CRASH assay with ere at *HMLa2.* Only the GFP channel is shown. These colonies occured when the founding cell of the colony has already experienced a ss-of-silencing event, so all daughter cells express GFP and not *RFP*. The *po/30-6* homozygote *(/) MATa/MATa* diploid (JRY11686) and the *po/30-79* homozygote <ii *MATa/MATa* diploid (JRY11688) both had extra-bright sectors and non-fluorescent sectors. No, or very few, sectors were observed in § *POL30 MATa* (JRY10790), *POL30 MATa/MATa* (JRY11159), **c** *POL30 matM MATa* (JRY11718), *po/30-6 MATa* (JRY11137), **c** *po/30-6 matMMATa* (JRY11719), *po/30-79 MATa* (JRY11141), or -*po/30-79 matMMATa* (JRY11720). **(B)** Patch-mating assay. Each cii indicated strain was patched onto complete medium plates seeded with a freshly-plated lawn of either *MATa* or *MATa* haploid cells with complementary auxotrophies. After −18 hours, mating patches were replica plated onto minimal medium plates. Growth occurs within the patch only if the indicated strains mated with the mating tester lawn.

Both homozygotes and hemizygotes of *pol30-6* and *pol30-79* had higher rates of spore inviability than expected whereas wild type and *pol30-8* did not (Table 2). The *pol30-6* and *pol30-79* alleles complemented one another by this assay, and their combination with *pol30-8* also reduced the high levels of spore inviability (Table 2). The consistency between spore inviability and recombination of the GFP cassette in *pol30-6* and *pol30-79* diploids suggested that the increased spore death might be a result of high levels of mitotic recombination. The inviability was suggestive of unequal or intra-chromosomal crossing over, since well-aligned reciprocal recombination would not be expected to cause inviability.

**Table 2.**
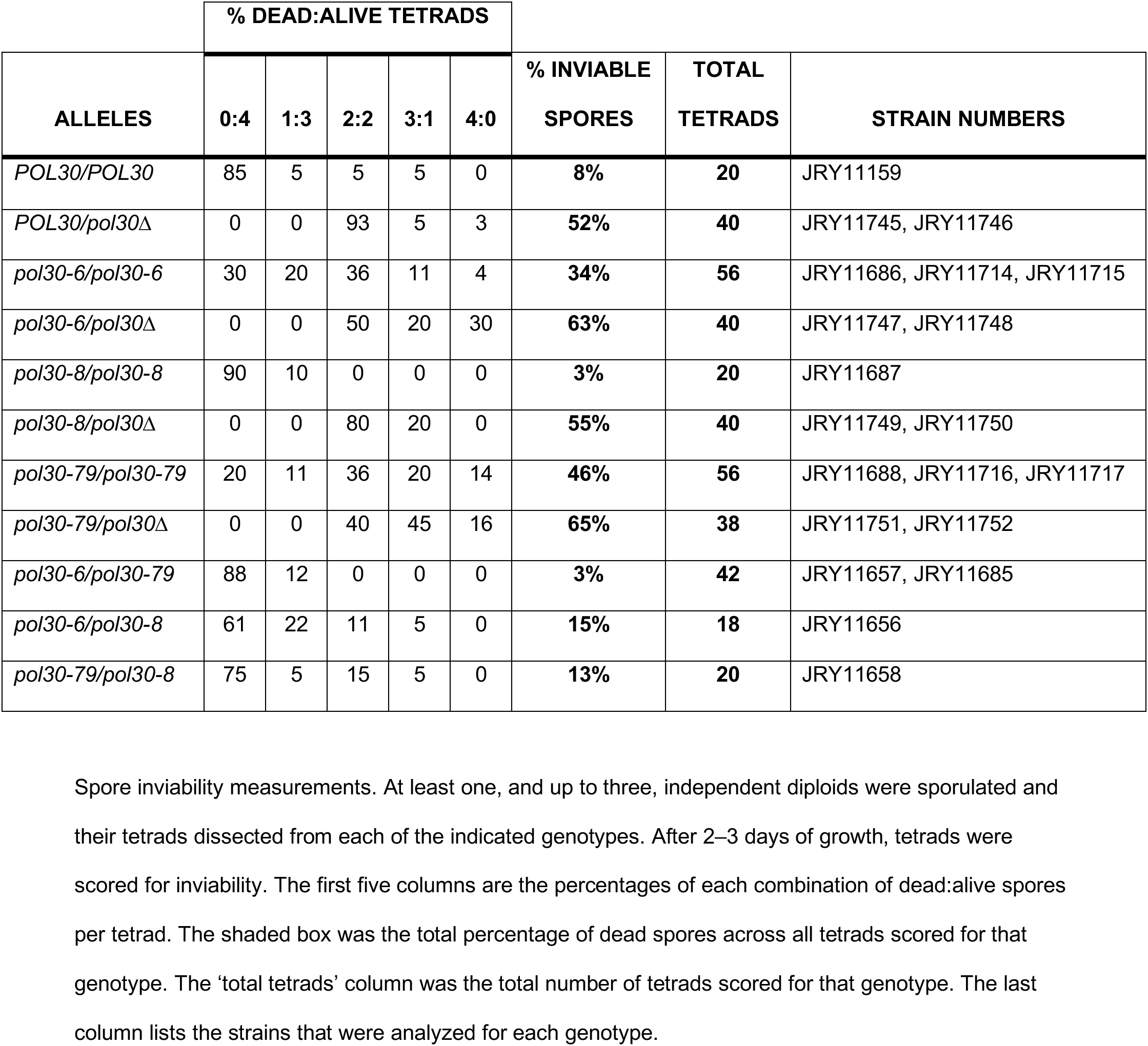
*pol30-6* and *pol30-79* cause high rates of mitotic recombination and gene conversion in diploids

Further evidence of genome instability of *pol30-6* and *pol30-79* mutants came from mating-type testing of diploids homozygous for these alleles. Diploid cells express both *MATa* and *MAT*α information, which prevents them from mating. However, if they undergo mitotic recombination between the centromere and *MAT* or gene conversion event at the *MAT* locus, they could become *MATa / MATa* or *MAT*α */ MAT*α, resulting in some cells in a patch of cells gaining the ability to mate with *MAT*α or *MATa* tester lawns, respectively. We patched each strain onto normal growth medium with a lawn of either *MATa* or *MAT*α haploids with complementary auxotrophies. After allowing time for mating, we replica plated these patches onto minimal media, selecting for diploid cells by the complementation of auxotrophic markers.

As expected, haploid *MATa* strains mated only with the *MAT*α tester (Figure 5B, top row). Additionally, *MAT*α */ mat*∆ diploids mated robustly with the *MATa* tester (Figure 5B, bottom row). There is some mitotic recombination/gene conversion that occurs in wild-type diploids allowing them to mate inefficiently with *MATa* and *MAT*α cells (Figure 5B, WT/WT MATa/α). However, *pol30-6* and *pol30-79* homozygous diploids had much higher levels of mitotic recombination/gene conversion, demonstrated by the greater density of colonies in those patches (Figure 5B, row 2).

Combination of *pol30-6* or *pol30-79* with a wild-type *POL30* allele or the *pol30-8* allele reduced the mating efficiency back to wild-type levels (Figure 5B, rows 3 and 4). Although still elevated compared to wild-type, *MAT*α */ mat*∆ diploids of *pol30-6* and *pol30-79* had lower amounts of mitotic recombination/gene conversion. Mutations that elevate the rate of chromosome loss would be expected to have a similar phenotype, but the involvement of mating type was suggestive of recombination events rather than chromosome losses being elevated in the homozygous diploids.

In contrast to the spore inviability results, where *pol30-6* and *pol30-79* complemented one another, there was no detectable complementation by this assay: The diploids with both *pol30-6* with *pol30-79* still had increased ability to mate as a diploid (Figure 5B, row 4) compared to the wild-type diploid.

### Coordination of histone chaperones at replication forks by PCNA was required for full transcriptional silencing

Because the *pol30* mutants appeared to have separable defects in heterochromatic silencing, we combined each of the alleles with known mutants affecting histone chaperone events at the replication fork: *cac1*∆*, dpb3*∆, and *mcm2-3A*. We made each combination in a CRASH assay strain and compared the sectoring phenotype of the double mutants with the corresponding single mutants.

*CAC1* is a subunit of the histone chaperone complex CAF-I. CAF-I deposits newly synthesized H3/H4 tetramers on daughter strands of DNA during replication (Smith and Stillman 1989, Serra-Cardona and Zhang 2018). Previous double-mutant analyses using a different silencing assay concluded that *pol30-8* results in loss of silencing through a defect in CAF-I activity, but that *pol30-6* and *pol30-79* act through a different mechanism (Zhang *et al.* 2000). In contrast to previous reports of a weak silencing defect for *cac1*∆ (Zhang *et al.* 2000; Huang *et al.* 2005), it had a severe sectoring phenotype in the CRASH assay, comparable to that of *pol30-8* alone (Figure 6A, 6B). The combination of *cac1*∆ with *pol30-8* was similar in phenotype to the single mutants, in agreement with previous results and the hypothesis that *pol30-8* and *cac1*∆ decrease silencing stability through the same mechanism (Figure 6A, 6B). Also, in agreement with previous results, the combination of *cac1*∆ with *pol30-6* or *pol30-79* worsened their phenotype significantly, suggesting that *pol30-6* and *pol30-79* had defects distinct from *cac1*∆ (Figure 6A, 6B).

**Figure 6.**
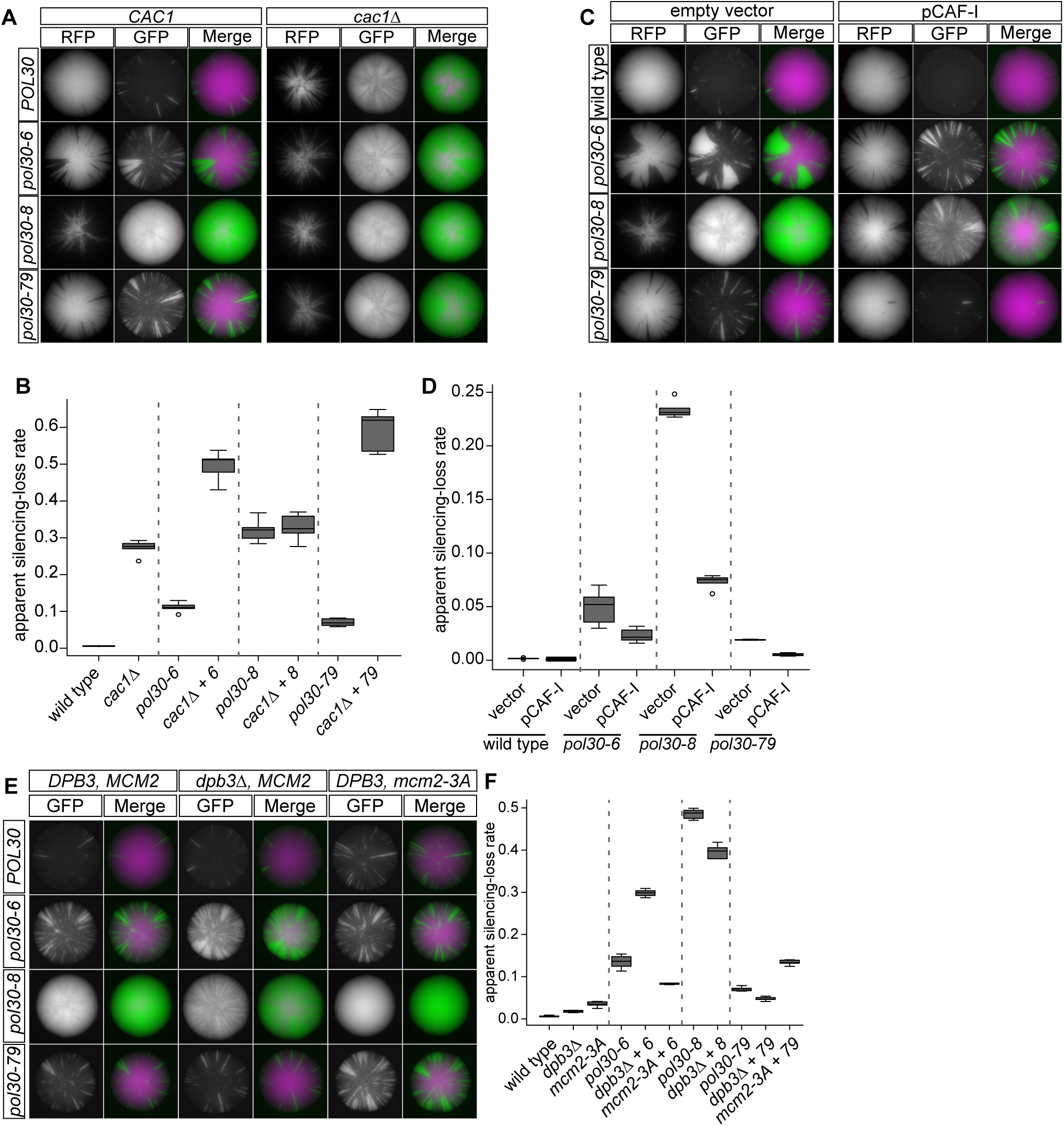
Coordination of histone chaperones by PCNA was required for transcriptional silencing. **(A)** Double-mutant analysis of *POL30* alleles with *cac1 A.* Representative images of colonies with *ere* at *HMLa2* in the CRASH assay. The left panel shows colonies with each of the *POL30* alleles with wild-type *CAC1* strain: *POL30* (JRY10790), *pol30-6* (JRY11137), *pol30-8* (JRY11188), -i— *pol30-79(*JRY11141). The right > panel shows colonies with each of the *POL30* alleles in combina¬tion with deletion of CAC1 *(cad* A): *POL30 cad A* (JRY11193), *pol30-6 cad A* (JRY11192), *pol30-8 cad A* (JRY11189), *pol30-79 cad A* (JRY11163) **(B)** The apparent silencing-loss rates for each of the strains in (A) were quantified by flow cytometry as described in Figure 1C **(C)** Overexpression of the CAF-I complex in combination with ■=■ *POL30* alleles. Representative images of colonies from the CRASH assay with *ere* at *HMLa* —i— 2. The left panel shows colonies •\9> with each of the *POL30* alleles in * combination with a 2p vector (pRS425): *POL30* (JRY11175), *pol30-6* (JRY11176), *pol30-8* (JRY11177), *po130-79* (JRY11178). The right panel shows colonies with each of the *POL30* alleles in combination with a 2g plasmid expressing all three subunits of the CAF-I complex, *CAC1, CAC2*, and *CAC3* (pJR3418): *POL30* pCAF-l (JRY11165), *pol30-6* pCAF-l (JRY11166), *pol30-8* pCAF-l (JRY11167), *pol30-79* pCAF-l (JRY11168). **(D)** The apparent silencing-loss rates for each of the strains in (C) were quantified by flow cytometry as described in Figure 1C. **(E)** Double-mutant analysis of *POL30* alleles with *dpb3A* and *mcm2-3A* alleles. Representative images of colonies from the CRASH assay with *ere* at *HMLa2.* In the left panel are each of the *POL30* alleles in a wild-type strain: *POL30* (JRY10790), *pol30-6* (JRY11137), *pol30-8* (JRY11188), *pol30-79* (JRY11141). In the middle panel are each of the *POL30* alleles in combination with deletion of DPB3 (*dpb3A)\ POL30 dpb3A* (JRY11760), *pol30-6 dpb3A* (JRY11806), *pol30-8 dpb3A* (JRY11808), *pol30-79 dpb3A* (JRY11810). In the right panel are each of the *POL30* alleles in combination with the *mcm2-3A allele. POL30 mcm2-3A* (JRY11812), *pol30-6 mcm2-3A* (JRY11987), *pol30-8 mcm2-3A* (JRY11989), *pol30-79 mcm2-3A* (JRY11991). **(F)** The apparent silencing-loss rates for each of the strains in (E) were quantified by flow cytometry as described in Figure 1C. The silencing-loss rate for *pol30-8 mcm2-3A* double mutant could not be quantified because it uniformly expressed GFP.

To further test each allele’s dependence on CAF-I for their silencing phenotype, we overexpressed the three subunits of CAF-I, *CAC1, CAC2*, and *CAC3*, from a 2μ plasmid (pCAF-I, pJR3418, Janke *et al.* 2018). If overexpression of CAF-I could suppress or rescue the phenotype of a *pol30* mutant, it would be strong evidence that the *pol30* mutant weakened silencing through reduced CAF-I activity. Compared to an empty vector control, overexpression of CAF-I strongly suppressed the phenotype of *pol30-8* (Figure 6C, 6D), in agreement with the conclusion from the double-mutant analysis between *pol30-8* and *cac1*∆ (Figure 6A, 6B). In contrast, overexpression of CAF-I in strains harboring *pol30-6* and *pol30-79* had no statistically significant effect on their silencing defects (Figure 6C, 6D).

Whereas CAF-I chaperones newly-synthesized histones, recent evidence implicates both Dpb3 and Mcm2 as having a role in chaperoning parental histones at the replication fork. Dpb3 and Dpb4 are part of DNA polymerase ε, the leading strand polymerase, and re-deposit parental (old) histones onto the leading strand during DNA replication (He *et al.* 2017; Bellelli *et al.* 2018). Mcm2 is a subunit of the replicative helicase MCM2-7 and is responsible for re-deposition of parental histones onto the lagging strand during DNA replication (Petryk *et al.* 2018; Gan *et al.* 2018). Because *MCM2* is an essential gene in yeast, we used the *mcm2-3A* allele, which has a defect in its histone chaperone activity but not in its helicase activity (Foltman *et al.* 2013).

In the CRASH assay, *dpb3*∆ and *mcm2-3A* on their own had minor but statistically insignificant sectoring phenotypes (Figure 6E, 6F). Their combination with different *pol30* mutants showed an interesting pattern: If combination of an allele with *dpb3*∆ enhanced the sectoring phenotype, then combination with *mcm2-3A* weakened it, and vice versa (Figure 6F). Except for the *pol30-6 dpb3*∆ combination, the differences in phenotype between the single and double mutants are modest, suggesting that these effects might be indirect.

## DISCUSSION

### The *pol30-8* allele does not cause epigenetically bi-stable states as was previously reported

Previous studies of transcriptional silencing in strains carrying *pol30-8* used the *ADE2* gene within the *HMR* locus as a reporter of gene silencing. When transcriptional silencing is disrupted, cells expressed *ADE2* and are white instead of red. *pol30-8* colonies have red sectors and white sectors, previously thought to represent two stable populations of cells: silencing *ADE2* or expressing *ADE2*. Surprisingly, even though *pol30-8* had highly unstable silencing in the CRASH assay, we found no evidence of bi-stable populations in strains carrying the *pol30-8* allele. The CRASH assay relies on alleles of *HML* and *HMR* that more closely resemble the native structure of those loci; it uses the native α*2* promoter and leaves the 5’ and 3’ UTRs and the α*1* gene intact (Dodson and Rine 2015), none of which is true for the previously used *HMR::ADE2* reporter. Moreover, the α-factor halo assay failed to reveal any clonally stable populations of cells expressing *HML* in any of the *pol30* mutants, nor did the *HML*α*2::RFP, HMR*α*2::GFP* reporter. Therefore, the *ADE2* sectors were likely an artifact of the metabolically responsive *ADE2* reporter in this context as we found no evidence of bi-stability among *pol30* mutants by two independent and metabolically neutral assays.

### Defective assembly versus maintenance of silenced chromatin *POL30* mutants

Although the expression of PCNA is highest in S phase and its major role in the cell occurs at replication forks, it is still present at lower levels in all stages of the cell cycle, functioning in DNA damage signaling and repair (Bauer and Burgers 1990). By monitoring loss-of-silencing events in unsynchronized cycling cells and cells arrested in G1 over time, we found that all three of the *pol30* mutants increased loss of silencing rates compared to wild type *POL30* only in cycling cells.

As the replication fork progresses, nucleosomes in front of the fork are disassembled and reassembled onto daughter strands and newly-synthesized nucleosomes are also deposited onto daughter strands. Sir proteins must reassemble on nucleosomes to re-set the heterochromatin state. Heterochromatin is then maintained through G2, M, and G1 until it is assembled again in the next round of replication. Since *pol30-6*, *pol30-8*, and *pol30-79* only caused loss of silencing in actively cycling cells, the predominant role of PCNA in silencing stability is most likely through heterochromatin assembly during S phase.

### Histone chaperones ensure the stability of heterochromatin through DNA replication

Previous results and supporting results in the CRASH assay shown here suggest that the unstable silencing caused by *pol30-8* is due to a defect in new histone deposition by the replication-coupled histone chaperone complex CAF-I (Zhang *et al.* 2000). Genetic evidence also suggests an interaction between two other histone chaperones, Hir1 and Asf1, and the *pol30-6* and *pol30-79* alleles (Sharp *et al.* 2001). If the *pol30* mutants caused slower or defective histone deposition during DNA replication, with fewer nucleosomes to bind, the SIR complex would be less able to properly block transcription at *HML* and *HMR* until the chromatin state was restored.

We explored the possibility that the *pol30* mutants might affect histone recycling from the mother DNA duplex to daughter strands chromatids by performing double-mutant analysis with *dpb3*∆ and *mcm2-3A*. The effects on CRASH sectoring phenotypes were minor but displayed an intriguing pattern: *mcm2-3A* and *dpb3*∆ had opposite effects for each given *pol30* mutant. These results could be interpreted as a leading strand or lagging strand bias for each allele. Biochemical studies of *pol30-79* show that it has a defect in binding to DNA polymerase δ (DNA Pol δ), the lagging-strand polymerase, but not DNA polymerase ε (DNA Pol ε), the leading-strand polymerase (Eissenberg *et al.* 1997). In contrast to *pol30-79, pol30-6* is completely unable to bind DNA Pol ε, with only reduced binding to DNA Pol δ (Ayyagari *et al.* 1995). Its CRASH phenotype in combination with *mcm2-3A* and *dpb3*∆ also mirror *pol30-79* analyses. Binding studies between the DNA polymerases and *pol30-8* have not been done, but the phenotype of the double-mutant analyses, interpreted in the light of a strand-bias model, suggested that *pol30-8* may exhibit weakened binding to DNA Pol δ but not Pol ε.

### A surprising effect of ploidy on silencing instability

Complementation tests revealed that any combination of *pol30-6, pol30-8*, or *pol30-79* in diploids complemented, resulting in colonies with fewer sectors in the CRASH assay than the haploids with each allele alone. These results suggested that *POL30* contributed at least two, and maybe three, separable roles in the assembly of stable silent chromatin. However, we also found that diploids carrying homozygous copies of *pol30-6* or *pol30-79* displayed no CRASH phenotype, and diploids homozygous for *pol30-8* had fewer sectors than a *pol30-8* haploid. Diploids homozygous for *POL30* alleles and expressing only *MAT*α mating type information all displayed the same suppression, establishing that the phenotype was not an unexpected manifestation of mating-type regulation. Likewise, hemizygosity of each allele in diploids failed to restore their CRASH sectoring phenotype back to haploids levels. Thus ploidy, independently of mating type and dosage of PCNA, changed the sensitivity of diploid cells to defects in histone deposition caused by the *pol30* mutant. To our knowledge, this wrinkle is unique among complementation tests, though we caution that most complementation tests are inadequately powered to detect the impact of ploidy. We note that none of the genes studied here are among the set shown to have ploidy-dependent impacts on their expression (Storchová *et al.* 2006)

*HML* and *HMR* cluster at the nuclear periphery with a higher local concentration of Sir proteins in the cluster than in the rest of the nucleoplasm (Gotta *et al.* 1996; Bystricky *et al.* 2009; Miele *et al.* 2009; Kirkland and Kamakaka 2013). The two copies of *HML* and *HMR* and the *SIR* genes in diploids might increase the local concentration of Sir proteins, despite the larger volume associated with increases in ploidy, enough that it could overcome a brief disruption in histone deposition during DNA replication in *pol30* mutants. Even though tethering to the nuclear periphery is not required for silencing in an otherwise wild-type strain (Gartenberg *et al.* 2004), the *pol30* mutants might create a sensitized background that is more dependent on clustering of the SIR complex with *HML* and *HMR* for maintenance of silencing through replication.

### A note on PCNA expression in hemizygotes

Compared to expression in homozygotes, the expression of PCNA in *pol30-6* and *pol30-79* strains did not decrease by half, whereas PCNA levels in *POL30* and *pol30-8* strains did. The expression of *POL30* is cell-cycle regulated (Bauer and Burgers 1990) and increases in response to DNA damage (Jelinsky and Samson 1999; Lee *et al.* 2007). Differences in cell-cycle distribution or levels of DNA damage in *pol30-6* and *pol30-79* hemizygotes compared to homozygotes would be compatible explanations for observations on PCNA levels produced by the various alleles.

### High levels of DNA damage and defective repair in *pol30-6* and *pol30-79*

The high rates of mitotic recombination and gene conversion we observed in *pol30-6* and *pol30-79* diploids presumably reflected higher levels of DNA damage in these mutants. Work from multiple labs shows that all three alleles, and especially *pol30-6* and *pol30-79*, are more sensitive to DNA damaging agents (Ayyagari *et al.* 1995; Eissenberg *et al.* 1997; Zhang *et al.* 2000; Miller *et al.* 2008). Additionally, *pol30-79* causes higher rates of substitutions and small insertions and deletions compared to wild-type cells (Eissenberg *et al.* 1997).

The published studies were done using haploid cells, where mitotic recombination is seldom detected. *MATa / MAT*α diploids avoid non-homologous end-joining because the a1-α2 transcription factor represses required genes *NEJ1* and *LIF1* (Astrom *et al.* 1999; Lee *et al.* 1999; Frank-Vaillant and Marcand 2001; Kegel *et al.* 2001; Valencia *et al.* 2001). Deleting *MATa* in the *pol30-6* and *pol30-79* homozygous diploids suppressed the increase in mitotic recombination, suggesting that the higher rates of mitotic recombination and gene conversion might be caused by homology-directed repair in lieu of non-homologous end-joining. We observed synthetic lethality in the haploid double mutants *pol30-6 rad54*∆*, pol30-79 rad54*∆, and *pol30-6 rad9*∆ and synthetic growth defects in the double mutants *pol30-79 rad9*∆*, pol30-6 H3K56R*, and *pol30-79 H3K56R* (data not shown). These synthetic phenotypes fit with a higher DNA damage load in strains carrying *pol30-6* and *pol30-79*. Our work in diploids provided more evidence that the *pol30-6* and *pol30-79* alleles have DNA damage and repair defects. It is unlikely that the mitotic recombination and silencing phenotypes of the *pol30-6* and *pol30-8* alleles are directly related since one phenotype was dependent on mating type (mitotic recombination) and the other was not (silencing).

The mutants *pol30-6* and *pol30-79* alleles reduce global levels of histone H3 lysine 56 (H3K56) acetylation (Recht *et al.* 2006; Miller *et al.* 2008), a histone modification that increases the affinity of CAF-I for H3/H4 dimers (Masumoto *et al.* 2005; Recht *et al.* 2006; Li *et al.* 2008). Misregulation of H3K56 acetylation, both hypoacetylation and hyperacetylation, is associated with increased DNA damage and sensitivity to DNA-damaging agents (Hyland *et al.* 2005; Masumoto *et al.* 2005; Recht *et al.* 2006; Clemente-Ruiz *et al.* 2011; Wurtele *et al.* 2012). These results could explain the high levels of mitotic recombination in *pol30-6* and *pol30-79*.

## ACKNOWLEDGEMENTS

We thank the Stillman and Burgers labs for plasmids carrying *POL30* alleles. We thank members of the Rine and Koshland laboratories for helpful discussions and Hector Nolla at the Cancer Research Laboratory Flow Cytometry Facility for technical help with flow cytometry. This work was supported by an NSF Predoctoral Fellowship (M.B.) and grants from the National Institutes of Health (GM031105 and GM120374 to J.R.).

