## Supplemental File 1 for "*POL30* alleles in *Saccharomyces cerevisiae* reveal complexities of the cell cycle and ploidy on heterochromatin assembly"

**FILE S1: SUPPLEMENTARY MATERIALS AND METHODS**

**Creation of *pol30-6* and *pol30-79* by URA3 swap:** The *pol30-6* (D41A, D42A) and *pol30-79* (L126A, I128A) alleles were created by integration of two overlapping gene blocks (gBlocks, Integrated DNA Technologies) at the *POL30* locus. Two gene blocks were used because the full sequence needed was too long to synthesize. One block contained the *pol30-6* or *pol30-79* allele followed by the first two-thirds of the *Candida albicans* *URA3* gene and the other contained the second two-thirds of the *Candida albicans URA3* gene followed by the *pol30-6* or *pol30-79* allele. The sequence of gene blocks is in Table S3. Selection for pop-out of the *URA3* gene was performed using the drug 5-fluoroorotic acid. Confirmation of the *pol30-6* and *pol30-8* allele was performed by PCR-amplification of the entire *POL30* locus followed by Sanger sequencing.

**Creation of *POL30* hemizygotes:** Hemizygotes were created by mating between *MATα* strains containing *POL30* (JRY11131), *pol30-6* (JRY11133), *pol30-8* (JRY11135), or *pol30-79* (JRY11137) with a *MATa* strain with a deletion of *POL30* (*pol30∆*) complemented by a plasmid carrying *POL30* with a *TRP1* marker (ZGY5-0, Zhang *et al.* 2000). The resulting diploids were streaked out for single colonies twice and checked for tryptophan auxotrophy, indicating they had lost the *POL30* plasmid. Hemizygosity for *POL30* was further confirmed by sporulation and tetrad analysis. Each strain resulted in at least 50% spore inviability (*POL30* is an essential gene) and no spores grew on CSM – Trp plates, indicating they did not carry the *POL30, TRP1* plasmid.

**Creation of tetraploid:** The tetraploid *POL30 / pol30∆ / pol30∆ / pol30∆* strain (JRY12026) was created in multiple steps. First, a *POL30* hemizygous diploid carrying a *POL30, TRP1* plasmid was screened for spontaneous *MATa/MATa* diploids by ability to mate with a haploid *MATα* mating tester. The resulting *MATa/MATa* diploid was confirmed by sporulation in the presence of nicotinamide and tetrad analysis; all spores mated only with a *MATα* mating tester. The remaining chromosomal copy of *POL30* in this strain was deleted and replaced with a *Kluyveromyces lactis LEU2* gene, leaving a *MATa/MATa* diploid with the only *POL30* gene present on a plasmid. This *MATa/MATa;* *pol30∆/pol30∆* diploid was mated with a *MATα/MATα* *POL30/pol30∆* hemizygous diploid (screened in the same manner as above for spontaneous *MATα/MATα* diploids). The resulting tetraploid was streaked out for single colonies twice and checked for tryptophan auxotrophy, indicating they had lost the *POL30* plasmid. The final strain, JRY12026, was confirmed by sporulation and tetrad dissection.
